# Bioinspired Design of Lysolytic Triterpenoid-peptide Conjugates that Kill African Trypanosomes

**DOI:** 10.1101/451252

**Authors:** W.-Matthias Leeder, Fabian Giehler, Juliane Joswig, H. Ulrich Göringer

**Affiliations:** Molecular Genetics, Darmstadt University of Technology, Schnittspahnstr. 10, 64287 Darmstadt, Germany; Helmholtz Zentrum München für Gesundheit und Umwelt (GmbH), Research Unit Gene Vectors, Munich, (Germany) and German Center for Infection Research (DZIF), Partner Site Munich, Munich, (Germany)

**Keywords:** African trypanosomes, sleeping sickness, trypanolytic factor (TLF), bioinspired design, synthetic TLF

## Abstract

Humans have evolved a natural immunity against Trypanosoma brucei infections, which is executed by two serum (lipo)protein complexes known as trypanolytic factors (TLF). Active TLF-ingredient is the primate-specific apolipoprotein L1 (ApoL1). The protein has a pore-forming activity that kills parasites by lysosomal and mitochondrial membrane fenestration. Of the many trypanosome subspecies only two are able to counteract the activity of ApoL1, which illustrates its evolutionary optimized design and trypanocidal potency. Here we ask the question whether a synthetic (syn)TLF can be synthesized using the design principles of the natural TLF-complexes but relying on different chemical building blocks. We demonstrate the stepwise development of triterpenoid-peptide conjugates, in which the triterpenoids act as a cell binding, uptake and lysosomal transport-moduls and the synthetic peptide GALA as a pH-sensitive, pore-forming lysolytic toxin. As designed, the conjugate kills infective-stage African trypanosomes through lysosomal lysis demonstrating proof-of-principle for the bioinspired, forward-design of a synTLF.

---

The discovery and development of new drugs and therapeutics is perhaps one of the most challenging tasks of translational research. In part due to the complexity of the process drug discovery is inherently slow and cost-and labor-intensive.^[1]^ While modern era drug discovery has considerably benefited from the large-scale compilation of genome-and proteome-data, from high-throughput screening (HTS) and combinatorial chemistry, the need for new concepts in drug discovery has not waned.^[2]^ A more recent addition to the field is the forward-design or re-engineering of drugs following the design principles of natural systems. The approach, also known as bioinspired design or biomimicry^[3]^ takes example from the powerful resource of biological systems that have evolved to respond and to adapt, which is the basis for self-healing, the immune response or natural resistance. By imitating the make-up and functionality of such systems and by reconstructing them using synthetic building blocks biomimetic drugs can be developed.^[4]^

The search for new therapeutics has been especially slow and arduous in the case of neglected tropical diseases.^[5]^ A prominent example for this is human African trypanosomiasis (HAT).^[6]^ HAT, also known as sleeping sickness, is caused by protozoan blood parasites of the genus *Trypanosoma.* The disease is fatal if untreated because trypanosomes evade the host immune response by repetitively altering their antigenic glycoprotein surface.^[7]^ This has thwarted all vaccine development efforts and as a consequence, HAT is treated by chemotherapy.^[8]^ Unfortunately, all of the five currently approved drugs (suramin, melarsoprol, pentamidine, eflornithine, nifurtimox) are unsatisfactory due to varying degrees of toxicity, low efficacies and the necessity for parenteral administration. Furthermore, drug resistance represents a growing problem especially in the case of the arsenical melarsoprol.^[9]^ As a consequence, the search for new and advanced HAT-therapeutics is urgent.^[6]^

Importantly, of the different trypanosome subspecies only *Trypanosoma brucei rhodesiense* and *Trypanosoma brucei gambiense* are causing HAT. All other trypanosome species are lysed in human serum due to the presence of an innate resistance in the form of two trypanolytic factors (TLF-1, TLF-2).^[10]^ Both are high molecular mass serum complexes that contain as a lysosome and mitochondrial membrane destroying activity the pore-forming, Bcl-2-like protein apolipoprotein L1 (ApoL1).^[11]^ TLF-1 binds to a specific receptor in the flagellar pocket (FP) of trypanosomes, which triggers its internalization and endosomal routing to the lysosome. Acidification activates the pore-forming activity of ApoL1 and allows the protein to permeate endosomal, lysosomal as well as mitochondrial membranes. This induces an influx of H_2_O and Cl^-^-ions, which results in the uncontrolled osmotic swelling of the lysosome^[12]^ in additon to the release of a mitochondrial endonuclease, which ultimately destroys the parasite (Figure 1A).^[13]^ Thus, ApoL1 represents a potent trypanocide and recombinant mutants as well as nanobody-coupled truncated versions of the protein have successfully been used as antitrypanocidals in mice.^[14]^ Unfortunately, the two HAT-causing trypanosome species are resistant to ApoL1. *Trypanosoma brucei rhodesiense* expresses the ApoL1-interacting protein SRA (serum resistance associated protein), which prevents ApoL1 from accumulating in the lysosome.^[15]^ *Trypanosoma brucei gambiense* relies on the parasite-specific glycoprotein TgsGP, which executes a membrane stiffening reaction, thus precluding the insertion of ApoL1 into membranes.^[16]^

**Figure 1.**
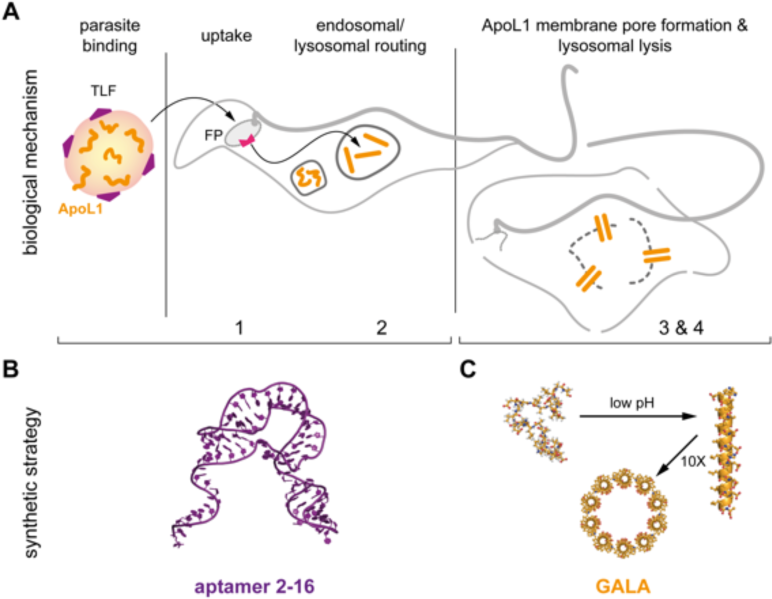
A) Mode-of-action of human TLF. Four stages can be distinguished: 1. Binding to the flagellar pocket (FP) of the parasite. 2. Uptake and endosomal routing to the lysosome. 3. pH-triggered pore formation of ApoL1. 4. Lysosomal membrane integration of ApoL1 followed by osmotic swelling and desintegration of the lysosome/cell. B) Synthetic strategy: A substitute for reaction steps 1 and 2 is the synthetic aptamer 2-16. C) Steps 3 and 4 can be executed by the synthetic, pore-forming peptide GALA.

Here we report the stepwise engineering of a bioinspired, synthetic trypanolytic factor (*syn*TLF). *syn*TLF emulates the make-up and mode of action of the natural TLF’s, however, it relies on different chemical compounds to potentially sidestep parasite resistance.

Starting point for the synthesis of a *syn*TLF was the selection of two synthetic molecules or modules that individually have been shown to execute some of the reaction steps of the natural TLF’s. As the “parasite-binding, uptake and lysosomal targeting”-modul we chose the trypanosome-specific RNA-aptamer 2-16 (Figure 1B).^[17]^ The synthetic RNA is 78nt long. It binds with high affinity (Kd 60nM) to the FP of the parasite after which it is internalized and routed to the lysosome.^[18]^ As a “pH-dependent, membrane disrupting”-modul we selected the synthetic peptide GALA (Figure 1C).^[19]^ GALA was originally designed to analyze the interaction of viral fusion proteins with membranes and consists of the 30 amino acid sequence WEAALAEALAEALAEHLAEALAEALEALAA. The glutamic acid residues within the sequence act as pH-sensing, protonateable groups and at neutral pH, the peptide is negatively charged adopting a random coil conformation. Shifting the pH to mild acidic conditions protonates the glutamic acids, thus forcing the peptide into an amphiphatic helix. In this conformation about 10 GALA-monomers multimerize to form an aqueous pore capable of penetrating lipid bilayers.^[20]^

Aptamer 2-16 was synthesized by “run-off” *in vitro* transcription and was purified by electrophoresis in denaturing, high resolution polyacrylamide gels. GALA was synthesized by solid-phase peptide synthesis and was purified by high performance liquid chromatography (HPLC). Site-specific coupling of the two reagents was achieved by reductive amination. For that the 3’-terminus of the aptamer was NaIO_4_-oxidized^[21]^ to generate 3’-terminal aldehyde groups, which at mild alkaline pH react with the N-terminal NH_2_-group of GALA. The resulting labile Schiff’s base was reduced to a stable secondary amide. After optimization of the reaction a maximal yield of 49% aptamer∼GALA conjugate was achieved. To test its trypanocidal activity, we incubated bloodstream-stage trypanosomes with the synthesized product up to a concentration of 10µM. The number of surviving cells was determined in a fluorescence-based life cell detection assay.^[22]^ As shown in Figure 2 aptamer∼GALA conjugates exhibit a dose-dependent cytotoxicity. At a concentration of 10µM ≥95% of the parasites are killed. This calculates to a lethal dose of 6×10^7^ aptamer∼GALA molecules per cell and a half-maximal lethal concentration (LC_50_) of 3.2±0.5µM (Figure 2). Importantly, neither the aptamer alone nor GALA alone show any inhibitory effect, even at concentrations up to 50µM (Figure 2). Thus, the cytoxicity is specific for the aptamer∼GALA conjugate.

**Figure 2.**
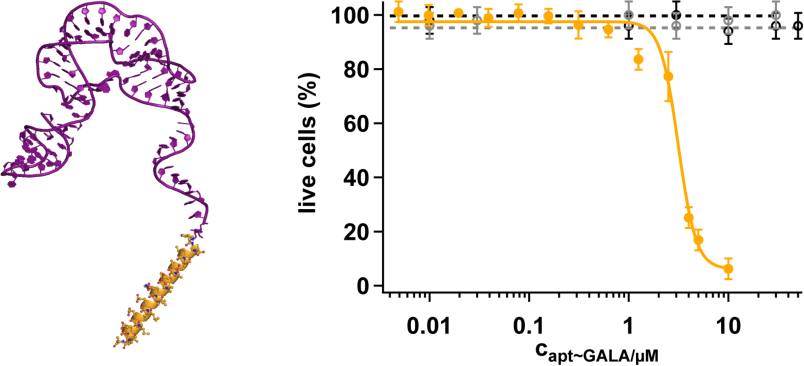
Left panel: 3D-model of aptamer 2-16∼GALA (aptamer: purple; GALA: yellow). Right panel: Dose-response curve of the aptamer∼GALA conjugate (filled circles, yellow). The half-maximal lethal concentration (LC_50_) calculates to 3.2±0.5µM. Non-conjugated 2-16 aptamer (grey open circles) and non-conjugated GALA (black open squares) show no cytotoxicity. Errors are standard deviations.

To further develop *syn*TLF, especially to reduce its molecular mass, we asked the question whether the aptamer domain of the conjugate can be substituted by a low molecular mass compound. For that we performed a high throughput (HT)-competition screen to identify molecules capable of competing with aptamer 2-16 for its binding site on the trypanosome surface.^[23]^ We used a compound library containing 1056 compounds with molecular masses ranging between 100-3500g/mol. About half of the compounds are synthetic molecules (50.3%), 37.4% are derived from plants and 15.1% from microbial species. 17.3% represent natural product derivatives and only 1% of the molecules are from animals (0.6%) or marine organisms (0.4%).

The library covers a diverse chemical space (Figure S1) and the screening experiment was characterized by a mean signal-to-background (S/B) ratio of 3.1 and a mean screening window coefficient (Z’) of 0.82.^[24]^ Using a cut-off of 3 standard deviations from control samples we identified 25 primary hits ranging in molecular mass between 234-1964g/mol. All hit-compounds were further analyzed in dose response curves, which identified six false-positives. The remaining 19 compounds (Tables S1,S2) cover a divers chemical space and include the plant-derived molecules Magnolol (C_18_H_48_O_2_) and Polygodial (C_15_H_22_O_2_), the microbial compounds Tyrocidine (C_66_H_87_N_13_O_13_) and Alamethicin (C_92_H_150_N_22_O_25_) and the synthetic (R,R)-Bis{2-[1-(methylamino)ethyl phenyl}diselenid (C_18_H_24_N_2_Se_2_). Importantly, five compounds represent substituted triterpenoids: Quercinic acid (C_31_H_50_O_4_), Nigranoic acid (C_30_H_46_O_4_), Aridanin (C_38_H_61_NO_8_), a highly substituted oleanic acid derivative (C_42_H_68_O_13_) and a plant compound with the molecular formula C_30_H_48_O_5_ (Figure 3A). This demonstrates that defined triterpenoid-type scaffolds can out-compete aptamer 2-16 from its cell surface binding site. Within this context it is important to note that trypanosomes are lipid auxotrophes. The parasites depend on the uptake of lipoproteins from the host serum, which includes the scavenging of sterols such as cholesterol esters and chlolesterol.^[25]^ As a consequence, we considered the identified triperpenoid-type scaffolds as promising candidates and used quercinic acid (QA) as a representative because of its favorable toxicity profile (all other compounds were found to be toxic at concentrations ≥30µg/mL). QA is a secondary metabolite derived from the fungi *Daedalia quercinia* and *Fomitopsis spraguei*.^[26]^ It consists of a lanost-8-en scaffold that is substituted with a C9-side chain terminating in a carboxyl-group (Figure 3A). To visualize the binding of QA to parasite cells and to confirm the aptamer-like uptake and intracellular routing to the lysosome we synthesized a biotin-substituted derivative of QA (QA_bio_). As shown in Figure S2, QA_bio_ binds to the FP of the parasite as expected. The molecule becomes internalized by endosomal uptake and is transported to the lysosome. As such, QA behaves identical to aptamer 2-16.^[18]^

**Figure 3.**
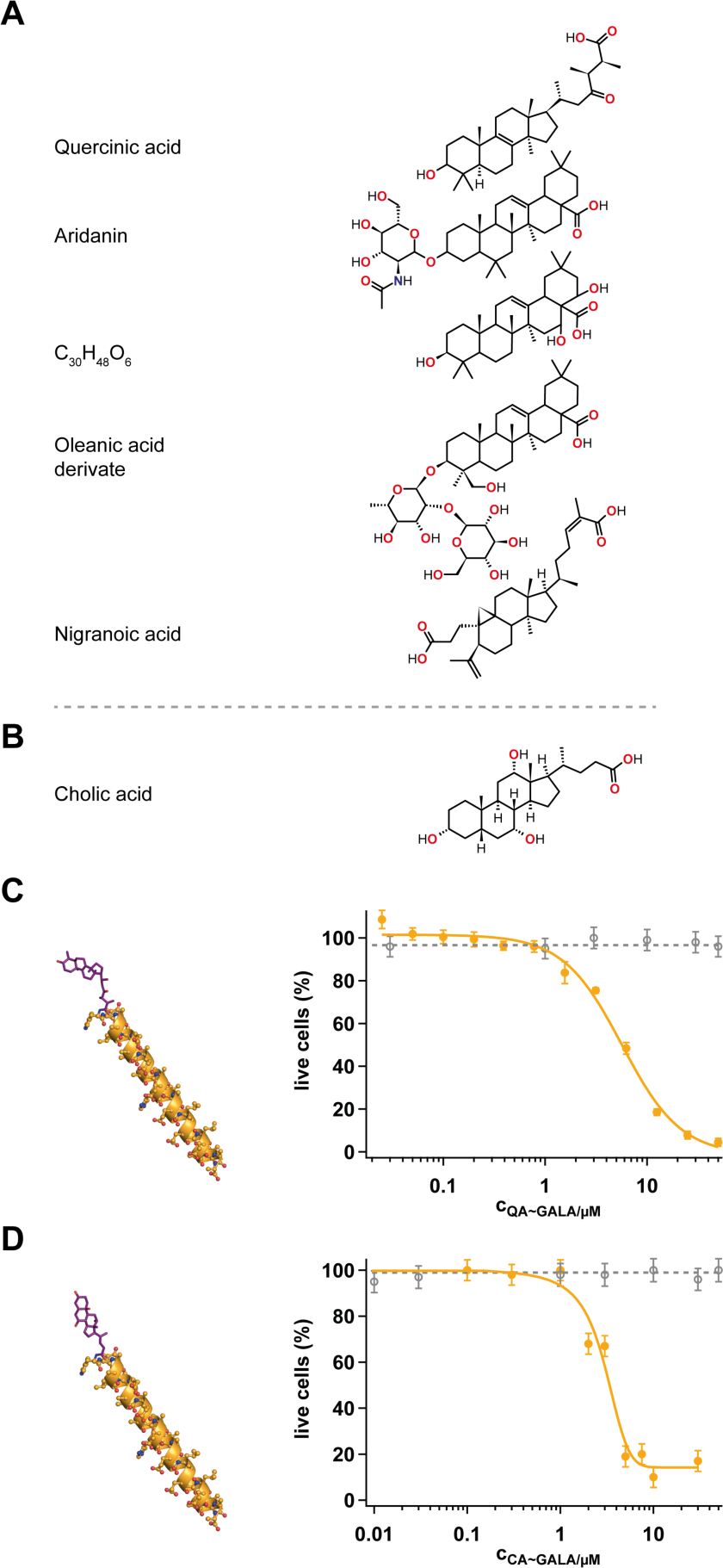
A) Structures of the triterpenoid-type hit compounds of the HT-aptamer displacement screen. B) Structure of cholic acid (CA), a compound not included in the library. C) Model and dose-response curve of QA∼GALA (filled circles, yellow). The LC_50_ calculates to 5.7±0.6µM. D) Model and dose-response plot of CA∼GALA (filled circles, yellow). The LC_50_ is at 2.9±0.4µM. Nonconjugated QA (grey, open circles in C) and non-conjugated CA (grey, open circles in D) show no cytotoxicity. Errors are standard deviations.

Next we synthesized QA∼GALA conjugates by covalently linking QA to the amino-terminus of the peptide using an EDC (1-Ethyl-3-(3-dimethylaminopropyl)carbodiimide)-based coupling reaction. QA∼GALA was synthesized with a yield ≥95% and its trypanolytic activity was analyzed as before. Figure 3C shows the corresponding dose-response curve. While QA alone has no effect, even at concentrations up to 50µM, QA∼GALA kills trypanosomes with an LC_50_ of 5.7±0.6µM almost identical to the aptamer∼GALA conjugate. Thus, the identification of QA as an aptamer 2-16 substitute reduced the molecular mass of *syn*TLF from 30.3kDa to 3.4kDa without any loss in function. Furthermore, we were able to demonstrate that next to QA other triterpenoid-type scaffolds can be used. As an example we tested cholic acid (CA, C_24_H_40_O_5_), a compound not included in the HTS-library (Figure 3B). CA was covalently attached to GALA using the same coupling chemistry as before and the CA∼GALA conjugate showed an LC_50_ of 2.9±0.4µM (Figure 3D), thus establishing the function of the triterpenoid-type scaffold.

Finally, we aimed at confirming the “designed” mode of action of *syn*TLF. Based on the biochemical properties of GALA, the peptide should adopt a helical structure upon entering the acidic interior of the lysosome, which in turn should activate the pore-forming activity of the peptide followed by the integration of the pores into the lysosomal membrane.^[27]^ An influx of Cl^-^-ions then should induce lysosomal swelling ultimately destroying the parasite. Figure 4A shows a representative experiment. Trypanosomes were treated with either CA∼GALA or QA∼GALA at concentrations 1.5-3.5-fold the LC_50_ and were microscopically monitored. Up to 60min the parasites show the typical elongated cell morphology of bloodstream-stage trypanosomes. At 75min the cells start to loose their slender shape showing signs of osmotic swelling and at 90min >75% of the parasites have a roundish, swollen phenotype. At 120min >95% of the cells are dead and a substantial amount of cells have desintegrated. Figure 4B shows a quantification of the process demonstrating the change in the length/width ratio of a parasite population treated with *syn*TLF. In order to verify that the *syn*TLF-induced osmotic swelling of the parasite cells is a consequence of the desintegration of the lysosome we pre-loaded the lysosomes of bloodstream-stage trypanosomes with fluorescently-labelled dextran as a fluid phase marker. Dextran pre-loaded cells were then incubated for 1.5h with either QA∼GALA or CA∼GALA at concentrations 1-2-fold the LC_50_. While untreated control cells showed no alterations of the integrity of the lysosome, *syn*TLF-treated cells are characterized by the loss of the punctate lysosomal staining (Figure 5) thus confirming the destruction of the lysosomal compartment as anticipated by the design of *syn*TLF.

**Figure 4.**
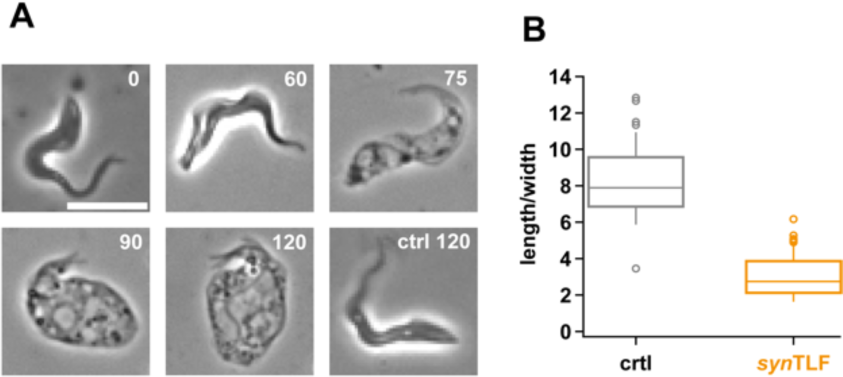
A) Kinetic of the *syn*TLF-driven osmotic swelling of bloodstream-stage trypanosomes. The experiment was performed with CA∼GALA at a concentration of 5µM (1.6-fold LC_50_). Numbers are incubation times in minutes. ctrl120: untreated parasites after 120min. Scale bar: 10µm. B) Box plot of the length/width ratio of *syn*TLF-treated trypanosomes in comparison to untreated control cells (ctrl).

**Figure 5.**
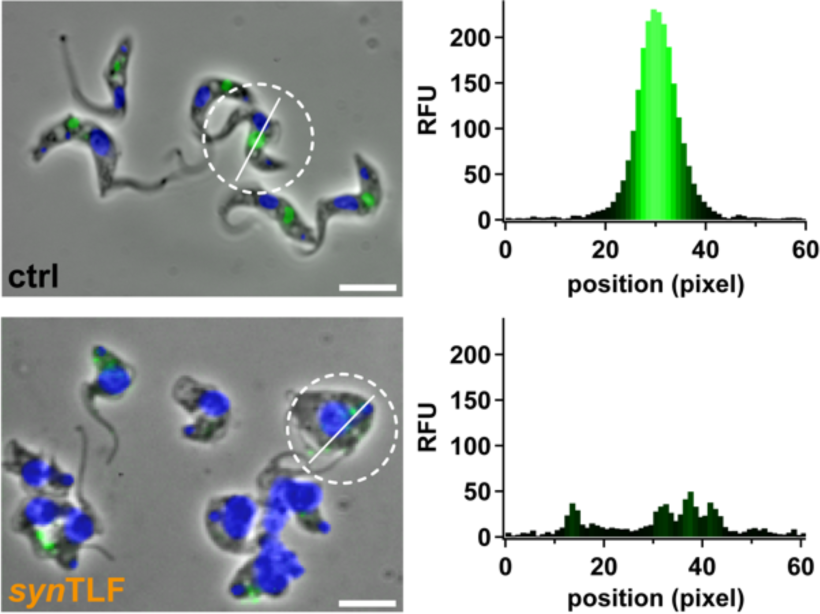
*syn*TLF-induced lysosomal lysis of bloodstream-stage trypanosomes. Lysosomes are pre-loaded with Alexa Fluor^®^ 488-labeled dextran (green). Nuclear-and kinetoplast-DNA are stained with Hoechst 33342 (blue). Top panel: Microscopic image of untreated trypanosomes (ctrl). Bottom panel: *syn*TLF-treated parasites (QA∼GALA, 5µM=0.9-fold LC_50_, 90min). Lysosomes loose their structural integrity after incubation with *syn*TLF, which was quantified by plotting the lysosomal staining profile in individual parasites (white line in dashed white circle). RFU: relative fluorescence unit. Scale bar: 10µm.

In summary, we have demonstrated proof-of-principle for the forward-design of a synthetic trypanolytic factor based on the design principles and mode-of-action of human TLF. We show that *syn*TLF is capable of killing bloodstream-stage African trypanosomes in a dose-dependent manner and that it acts by destroying the lysosomal compartment of the parasite as projected. This illustrates the potential of the bioinspired design approach. Since *syn*TLF relies on different chemical building blocks than its natural counterpart this raises the possibility that the mechanism by which HAT-causing trypanosomes bypass the natural trypanolytic activity, can be circumvented. In addition, replacing ApoL1 with GALA likely sidesteps the problem that certain ApoL1-variants are associated with an increased risk of kidney disease.^[28]^ Lastly, we foresee the design of fully synthetic TLF’s for instance by using non-natural triterpenoids,^[29]^ together with other pH-responsive, membrane-disturbing molecules,^[30]^ with perhaps even higher trypanocidal potencies.

## Acknowledgements

We thank R. Knieß and M. Brecht for experimental input and A. Adler for early work on the project. M. Homann is thanked for discussions and E. Kruse and C. Del Campo for critically reading the manuscript. The work was supported by the Hessian Ministry for Science and Art through the LOEWE-project CompuGene, the Deutsche Forschungsgemeinschaft (SFB902) and the Dr. Illing-Foundation for Molecular Chemistry. The Leibniz-Institute for Natural Product Research and Infection Biology in Jena is acknowledged for providing the HTS-compound library.

## Supporting Information

### 1 SI Methods

#### 1.1 Trypanosome cell growth

The bloodstream life cycle stage of *Trypanosome brucei brucei* strain 427 (MITat serodeme, variant clones MITat 1.2 and MITat 1.4) (Cross, 1975) was grown in HMI-9 medium (Hirumi and Hirumi, 1994) supplemented with 10% (v/v) heat-inactivated fetal calf serum (FCS). Parasites were grown at 37°C in 95% air, 5% CO_2_. Parasite cell densities were determined by automated cell counting.

#### 1.2 Aptamer and peptide synthesis

RNA-aptamer 2-16 (78nt) was synthesized by run-off transcription following standard procdures. The full-length RNA was separated from truncated reaction products by electrophoresis in 8M urea-containing 10% (w/v) polyacrylamide gels and passively eluted from the gel-matrix. (^32^P)-labelled aptamer preparations were generated using α-(^32^P)-ATP in the transcription mix. Aptamer concentrations were determined by UV absorbancy measurements at 260nm using a molar extinction coefficient (ε_260_) of 700900 M^−1^ cm^−1^. Oxidation of RNA 3’-ends (≤0.3mg) was performed in 50mM NaOAc pH4.5, 100mM NaCl, 10mM MgCl_2_ containing 8.3mM NaIO_4_ (Göringer et al., 1984). After incubation for 16h at 4°C in the dark the oxidized RNA was desalted by size exclusion chromatography followed by EtOH precipitation. The completeness of the reaction was tested by 5’-(^32^P)-pCp ligation using T4 RNA-ligase. GALA (WEAALAEALAEALAEHLAEALAEALEALAA) was synthesized by solid phase synthesis (Eurogentec), purified by reversed phase HPLC and verified by mass spectrometry.

#### 1.3 Synthesis of aptamer∼GALA conjugates

GALA was covalently coupled to aptamer 2-16 by reductive amination. The 3’-oxidized RNA (2µM) was dissolved in phosphate-buffered saline (PBS): 20mM Na_x_H_y_PO_4_ pH 7.4, 120mM NaCl, 5mM KCl, 2mM MgCl_2_ and incubated with 1-2mM GALA in the presence of 0.5– 1M KCl. NaCNBH_3_ (50mM) was added and the reaction was further incubated for 14-16h at room temperature (RT). Unreacted GALA was removed by size exclusion chromatography and samples were analyzed in denaturing polyacrylamid gels followed by SybrGreen I staining. Purification of the peptide/aptamer-conjugate from unreacted RNA was performed by electrophoresis in 2% (w/v) agarose gels. The conjugate was cut from the gel and passively eluted.

#### 1.4 Synthesis of QA∼ and CA∼GALA conjugates

Quercinic acid (QA, C_31_H_50_O_4_) was covalently linked to the amino-terminus of GALA following an EDC (1-Ethyl-3-(3-dimethylamino propyl)carbodiimide)-mediated protocol. The coupling was performed in 0.05mL using 1mM QA in dimethyl sulfoxide (DMSO) and 0.5mM GALA in 0.1M 3-morpholinopropane-1-sulfonic acid (MOPS) pH7 in the presence of 10mM EDC. The reaction was performed at RT for 3h with a yield >95%. Cholic acid (CA, C_24_H_40_O_5_) was coupled similarly following a two-step reaction. For that 0.5mmol CA, 0.35mmol *N*-hydroxysulfosuccinimide (sulfo-NHS) and 0.87mmol EDC were mixed in 20mL anhydrous tetrahydrofuran (THF) and incubated at 4°C over night. The solution was filtered and the filtrate poured into cold n-hexane. The succinimido-CA precipitate was dried and analyzed by ESI-mass spectrometry. The GALA-conjugation was performed using 0.6mM succinimidocholate and 0.3mM GALA in 25mM 2-(N-morpholino)ethanesulfonic acid (MES) pH6.5 for 24h at 4°C resulting in a 75% yield (Sehgal and Vijay 1994). Both conjugates were gelelectrophoretically analyzed in Tricine-SDS (sodium dodecyl sulfate) gels (Schägger, 2006).

#### 1.5 Synthesis of biotinylated aptamer 2-16 and biotinylated QA

The synthesis af biotinylated QA (QA_bio_) was performed in 0.03mL 0.1M MES pH5 containing 0.1mM QA, 17mM pentylamine-biotin (ThermoFisher Scientific) and 170mM EDC. Incubation was for 2h at RT. Aptamer 2-16 was biotinylated co-transcriptionally using 10µM biotin-16-UTP (Sigma-Aldrich) in the transcription mix.

#### 1.6 Cytotoxicity measurements

The cytotoxicity of GALA, all GALA-conjugates and all primary HTS-hit compounds was analyzed in a fluorescence-based live cell detection assay using Calcein *O*,*O*’-diacetate tetrakis(acetoxy methyl) ester (Calcein-AM, λex 496nm, λ_em_ 516nm) (Lichtenfels et al., 1994). Infective-stage trypanosomes (10^8^ cells/mL) were resuspended in PBSG (PBS, 20mM glucose) and 0.01-0.1mL of the cell suspension were incubated in the presence/absence of varying concentrations of the different compounds for 30min at RT. Samples were mixed with an equal volume of 5µM Calcein-AM in PBSG and incubated for 1h at RT in the dark. The fluorescence at 516nm was recorded to derive dose-response curves for the determination of LC_50-_values. Data points were measured in triplicate and fitted to a sigmoidal function. Errors are standard deviations.

#### 1.7 Fluorescence *in situ* hybridization and lysosomal integrity analysis

Trypanosomes (2×10^8^ cells/mL) in 0.1mL cold PBSG were incubated with QA_bio_ (33-165ng) for 30-60min at 4°C. Cells were washed, resuspended in 0.1mL PBSG and bound QA_bio_ was detected using a monoclonal Cy™3-conjugated anti-biotin antibody (dianova). Lysosomes were visualized by using Alexa Fluor^®^ 488-conjugated dextran (10kDa, Molecular Probes) as a fluid phase marker. Trypanosomes (10^7^ cells) were resuspended in PBS containing 5mg/mL Alexa Fluor^®^ 488 dextran. After incubation for 15min at RT in the dark, cells were washed and incubated in the presence of 5-10µM of the different GALA-conjugates. After 30, 60, 90, 120min of incubation, cells were fixed in 4% (w/v) paraformaldehyde and analyzed by fluorescence microscopy (Zeiss Axioskop2, Zeiss Plan-NEOFLUAR 100x objective). Nuclear-and kinetoplast-DNA was stained with Hoechst 33342. Digital images were captured (Zeiss AxioCam MRm) and processed using IPlab 3.6 (Scanalytics) and AxioVision 4.8.1.0 (Zeiss).

#### 1.8 High throughput (HT) competition screen

The compound library was kindly provided by the Leibniz-Institute for Natural Product Research and Infection Biology (HKI), Jena, Germany. It contains 1056 compounds derived from plants (37.4%), microbial (15.1%), natural products (17.3%), animals (0.6%) and marine organisms (0.4%). Molecular masses vary between 100-3500g/mol. A summary of the chemical nature of the different compounds is detailed in Figure S1. All compounds were dissolved in DMSO at a concentration of 1mg/mL. For the HT-competition screen 2×10^7^ bloodstream stage trypanosomes in 0.1mL PBSG, 0.1µg/µL BSA, 0.1µg/µL yeast tRNA were incubated with 1-10fmol (^32^P)-labelled RNA aptamer 2-16 in the presence of 3µg of each library compound and incubated for 20min. The reaction was stopped by the addition of 0.1mL dibutyl phthalate followed by centrifugation for 1min at 21000xg (Homann et al. 2006; Wille et al., 1998). Samples were shock frozen in liquid N_2_, the cell pellet-containing tip of the vial was cut off and scintillation counted to determine the percentage of cell-bound (^32^P)-labelled aptamer. An excess of non-radioactive 2-16 RNA served as positive control and DMSO-only samples were used a a negative controls. The quality of the HT-screen was determined by evaluating the signal-to-background (S/B) ratio: S/B=µsignal/µbackground and the mean screening window coefficient (Z’): Z’=1-(3∂_control+_ + 3∂_control-_) / |µ_control+_-µ_control-_| (µ=mean, ∂=standard deviation) (Zhang et al., 1999).

## 2 SI Figures and Tables

**Figure S1.**
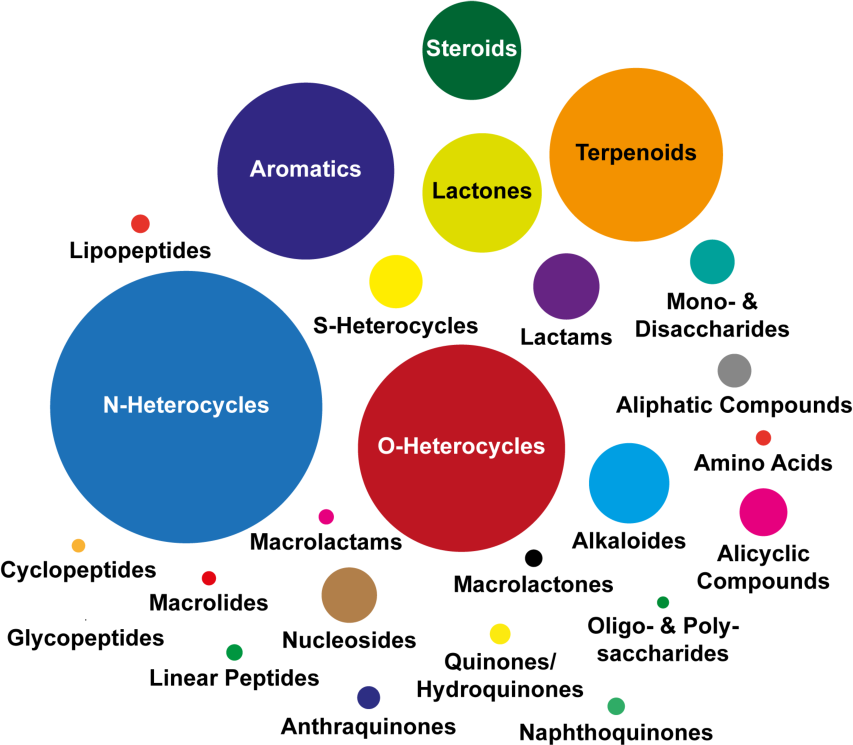
Molecular diversity of the HTS-compound library. The different chemical classes are shown as colored circles with diameters reflecting the number of molecules in the library.

**Figure S2.**
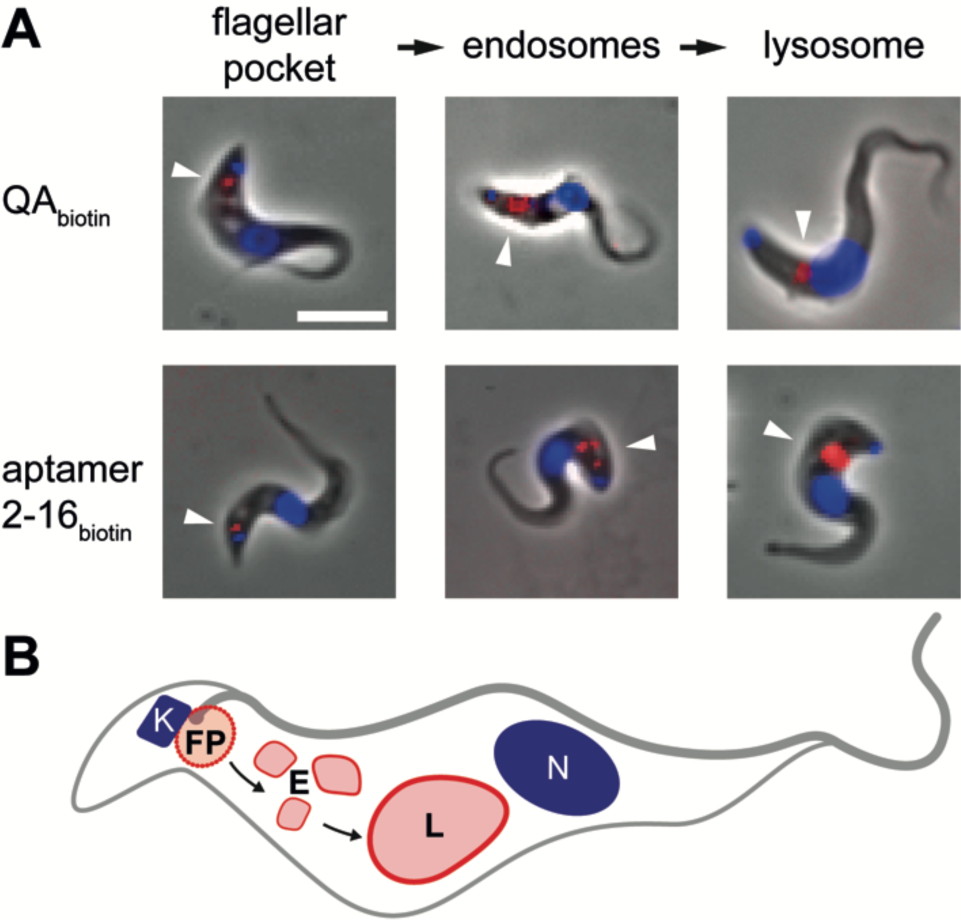
A) Microscopic analysis of the binding and fate of biotinylated quercinic acid (QA_biotin_) to bloodstream-stage trypanosomes in comparison to biotinylated preparations of aptamer 2-16 (aptamer 2-16_biotin_). Initial binding to the flagellar pocket is followed by endosomal uptake and transport to the lysosome. QA_biotin_ behaves identical to aptamer 2-16. B) Sketch of a trypanosome cell emphasizing the binding/transport pathway. Nuclear DNA (N), kinetoplast DNA (K), flagellar pocket (FP), endosomes (E), lysosome (L). Scale bar 10µm.

**Table S1.**
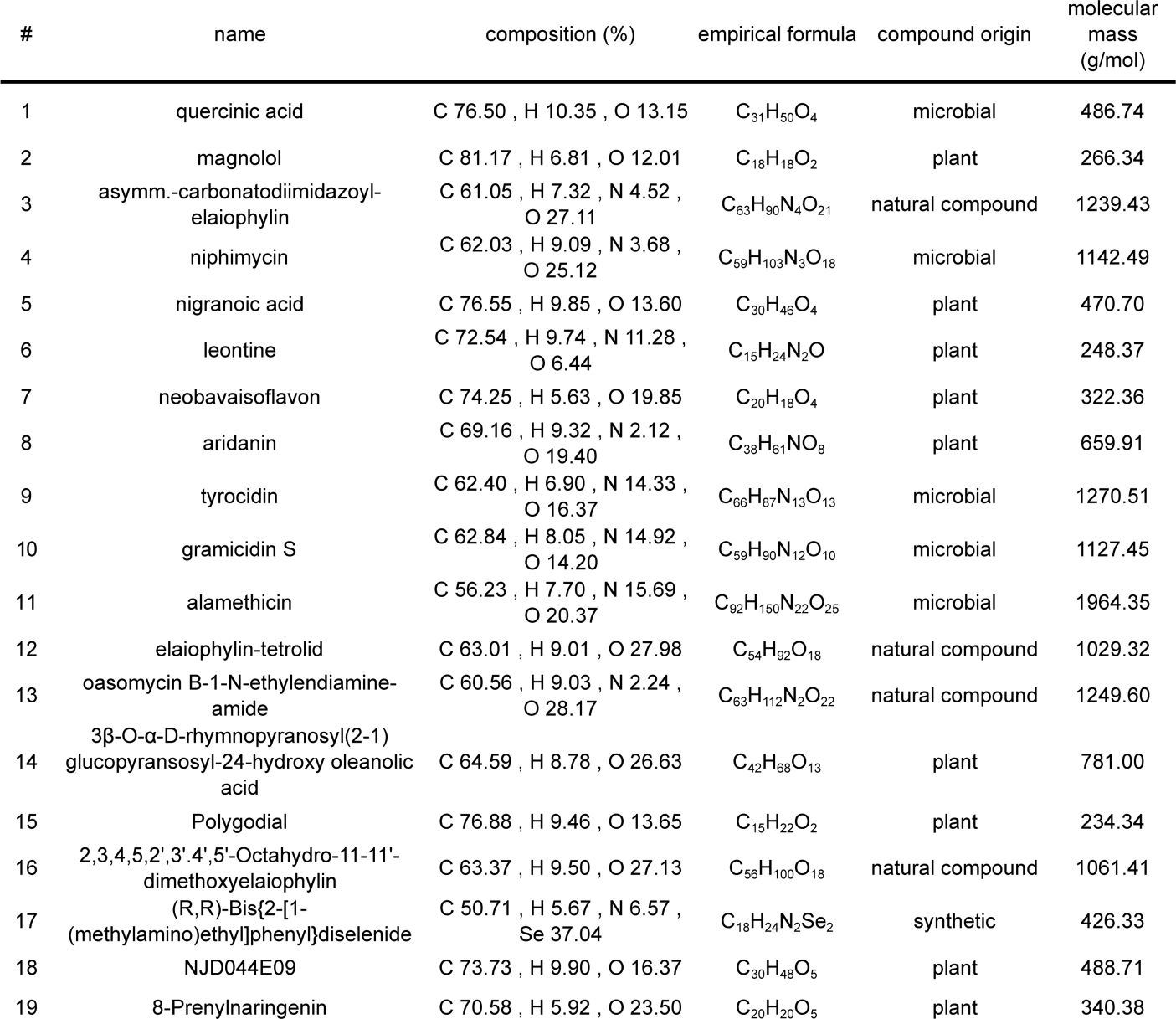
Primary hit compounds of the HT-aptamer competion screen.

**Table S2.**
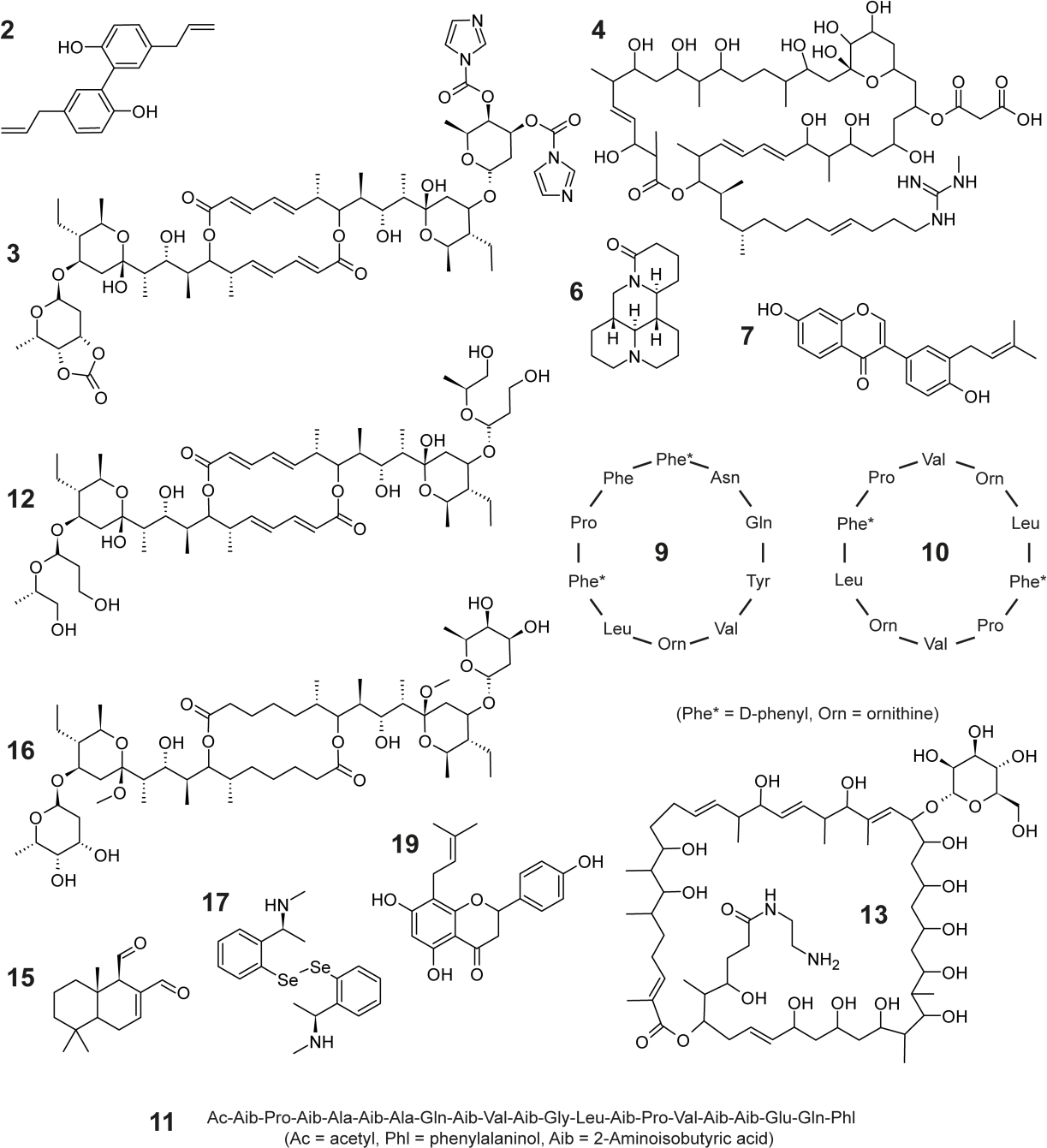
Hit compound structures of the HT-aptamer competion screen. The structures of the triterpenoid compounds 1, 5, 8, 14 and 18 (Table S1) are given in Figure 3A.

## References

[1] J. Eder, P. L. Herrling, Handb. Exp. Pharmacol. 2016, 232, 3–22.

[2] J. Boström, D. G. Brown, R. J. Young, G. M. Keserü, Nat. Rev. Drug Discov. 2018, doi: 10.1038/nrd.2018.116. [Epub ahead of print]

[3] R. Raman, R. Bashir, Adv. Healthc. Mater. 2017, 6, 1700496.

[4] a) S. Xue, J. Yin, J. Shao, Y. Yu, L. Yang, Y. Wang, M. Xie, M. Fussenegger, H. Ye, Mol. Ther. 2017, 25, 443–455; b) F. Rabanal, A. Grau-Campistany, X. Vila-Farrés, J. Gonzalez-Linares, M. Borràs, J. Vila, A. Manresa, Y. Cajal, Scientific Rep. 2015, 5, 10558.

[5] M. De Rycker, B. Baragaña, S. L. Duce, I. H. Gilbert, Nature 2018, 559, 498–506.

[6] a) M. C. Field, D. Horn, A. H. Fairlamb, M. A. Ferguson, D. W. Gray, K. D. Read, M. De Rycker, L. S. Torrie, P. G. Wyatt, S. Wyllie, I. H. Gilbert, Nat. Rev. Microbiol. 2017, 15, 217–231; b) C. H. Baker, S. C. Welburn, Trends Parasitol. 2018, pii: S1471–4922(18)30171–5.

[7] P. T. Manna, C. Boehm, K. F. Leung, S. K. Natesan, M. C. Field, Trends Parasitol. 2014, 30, 251–258.

[8] M. P. Barrett, S. L. Croft, Br. Med. Bull. 2012, 104, 175–196.

[9] A. H. Fairlamb, D. Horn, Trends Parasitol. 2018, 34, 481–492.

[10] B. Vanhollebeke, E. Pays, Mol. Microbiol. 2010, 76, 806–814.

[11] L. Vanhamme, F. Paturiaux-Hanocq, P. Poelvoorde, D. P. Nolan, L. Lins, J. Van Den Abbeele, A. Pays, P. Tebabi, H. Van Xong, A. Jacquet, N. Moguilevsky, M. Dieu, J. P. Kane, P. De Baetselier, R. Brasseur, E. Pays, Nature 2003, 422, 83–87.

[12] a) D. Pérez-Morga, B. Vanhollebeke, F. Paturiaux-Hanocq, D. P. Nolan, L. Lins, F. Homblé, L. Vanhamme, P. Tebabi, A. Pays, P. Poelvoorde, A. Jacquet, R. Brasseur, E. Pays, Science 2005, 309, 469–472; b) J. Bruno, N. Pozzi, J. Oliva, J. C. Edwards, J. Biol. Chem. 2017, 292, 18344–18353.

[13] G. Vanwalleghem, F. Fontaine, L. Lecordier, P. Tebabi, K. Klewe, D. P. Nolan, Y. Yamaryo-Botté, C. Botté, A. Kremer, G. S. Burkard, J. Rassow, I. Roditi, D. Pérez-Morga, E. Pays, Nat. Commun. 2015, 6, 8078.

[14] a) T. N. Baral, S. Magez, B. Stijlemans, K. Conrath, B. Vanhollebeke, E. Pays, S. Muyldermans, P. De Baetselier, Nat. Med. 2006, 12, 580–584; b) F. Fontaine, L. Lecordier, G. Vanwalleghem, P. Uzureau, N. Van Reet, M. Fontaine, P. Tebabi, B. Vanhollebeke, P. Büscher, D. Pérez-Morga, E. Pays, Nat. Microbiol. 2017, 2, 1500–1506.

[15] M. W. Oli, L. F. Cotlin, A. M. Shiflett, S. L. Hajduk, Eukaryot. Cell 2006, 5, 132–139.

[16] P. Uzureau, S. Uzureau, L. Lecordier, F. Fontaine, P. Tebabi, F. Homblé, A. Grélard, V. Zhendre, D. P. Nolan, L. Lins, J. M. Crowet, A. Pays, C. Felu, P. Poelvoorde, B. Vanhollebeke, S. K. Moestrup, J. Lyngsø, J. S. Pedersen, J. C. Mottram, E. J. Dufourc, D. Pérez-Morga, E. Pays, Nature 2013, 501, 430–434.

[17] M. Homann, H. U. Göringer, Nucleic Acids Res. 1999, 27, 2006–2014.

[18] a) M. Homann, H. U. Göringer, Bioorg. Med. Chem. 2001, 9, 2571–2580; b) A. Adler, N. Forster, M. Homann, H. U. Göringer, Comb. Chem. High Throughput Screen. 2008, 11, 16–23.

[19] a) N. K. Subbarao, R. A. Parente, F. C. Szoka Jr., L. Nadasdi, K. Pongracz, Biochemistry 1987, 26, 2964–2972; b) R. A. Parente, S. Nir, F. C. SzokaJr., J. Biol. Chem. 1988, 263, 4724–4730.

[20] W. Li, F. Nicol, F. C. SzokaJr., Adv. Drug Delivery Rev. 2004, 56, 967–985.

[21] H. U. Göringer, S. Bertram, R. Wagner, J. Biol. Chem. 1984, 259, 491–496.

[22] R. Lichtenfels, W. E. Biddison, H. Schulz, A. B. Vogt, R. Martin, J. Immunol. Methods 1994, 172, 227–239.

[23] a) L. S. Green, C. Bell, N. Janjic, Biotechniques 2001, 30, 1094–1096; b) P. Burgstaller, A. Girod, M. Blind, Drug Discov. Today 2002, 7, 1221–1228.

[24] J. H. Zhang, T. D. Chung, K. R. Oldenburg, J. Biomol. Screen. 1999, 4, 67–73.

[25] H. P. Green, M. Del Pilar Molina Portela, E. N. St. Jean, E. B. Lugli, J. Raper, J. Biol. Chem. 2003, 278, 422–427.

[26] a) J. Rösecke, W. A. König, Phytochem. 2000, 54, 757–762; b) D. N. Quang, Y. Arakawa, T. Hashimoto, Y. Asakawa, Phytochem. 2005, 66, 1656–1661.

[27] H. S. Choi, J. Huh, W. H. Jo, Biomacromol. 2006, 7, 403–406.

[28] A. K. Sharma, D. J. Friedman, M. R. Pollak, S. L. Alper SL. FEBS J. 2016, 283, 1846–1862.

[29] M. B. Sporn, K. T. Liby, M. M. Yore, L. Fu, J. M. Lopchuk, G. W. Gribble, J. Nat. Prod. 2011, 74, 537–545.

[30] A. Erazo-Oliveras, K. Najjar, L. Dayani, T. Y. Wang, G. A. Johnson, J. P. Pellois, Nat. Methods 2014, 11, 861–867.

## SI References

G. A. Cross, Parasitology 1975, 71, 393–417.

H. U. Göringer, S. Bertram, R. Wagner, J. Biol. Chem. 1984, 259, 491–496.

H. Hirumi, K. Hirumi, Parasitol Today. 1994, 10, 80–84.

M. Homann, M. Lorger, M. Engstler, M. Zacharias, H. U. Göringer, Comb. Chem. High Throughput Screen. 2006, 9, 491–499.

R. Lichtenfels, W. E. Biddison, H. Schulz, A. B. Vogt, R. Martin, J. Immunol. Methods 1994 172, 227–239.

H. Schägger, Nat. Protoc. 2006, 1, 16–22.

D. Sehgal, I. K. Vijay, Anal. Biochem. 1994, 218, 87–91.

U. Wille, B. Schade, M. Duszenko, Eur. J. Biochem. 1998, 256, 245–250.

J. H. Zhang, T. D. Chung, K. R. Oldenburg, J. Biomol. Screen. 1999, 4, 67–73.

